# Verification of *Arabidopsis* stock collections using SNPmatch - an algorithm for genotyping high-plexed samples

**DOI:** 10.1101/109520

**Authors:** Rahul Pisupati, Ilka Reichardt, Ümit Seren, Pamela Korte, Viktoria Nizhynska, Envel Kerdaffrec, Kristina Uzunova, Fernando Rabanal, Daniele Filiault, Magnus Nordborg

## Abstract

Large-scale studies such as the *Arabidopsis thaliana* 1001 Genomes Project aim to understand genetic variation in populations and link it to phenotypic variation. Such studies require routine genotyping of stocks to avoid sample contamination and mix-ups. To genotype samples efficiently and economically, sequencing must be inexpensive and data processing simple. Here we present SNPmatch, a tool which identifies the most likely strain (inbred line, or “accession”) from a SNP database. We tested the tool by performing low-coverage sequencing of over 2000 strains. SNPmatch could readily genotype samples correctly from 1-fold coverage sequencing data, and could also identify the parents of F1 or F2 individuals. SNPmatch can be run either on the command line or through AraGeno (<u>https://arageno.gmi.oeaw.ac.at</u>), a web interface that permits sample genotyping from a user-uploaded VCF or BED file.

Availability and implementation: https://github.com/Gregor-Mendel-Institute/SNPmatch.git

## Background & Summary

Sample contamination is an unavoidable problem when large-scale experiments are performed. For example, cell lines are frequently misidentified or contaminated, and the need for validation has long been underappreciated^1,2^. The same problem applies to any germplasm collection. With an increasing number of experiments utilizing natural variation, plant seed resources are growing rapidly. In *Arabidopsis thaliana*, the genomes of 1135 strains have recently been sequenced^3^ and this panel is now widely used. The need for verifying seed stock is clear^4^. Routine quality checks of seed stock genotypes can guard against common mistakes such as tube mislabeling, seed contamination during harvesting, or sample mix ups. In principle, genotyping can easily be performed by short-read sequencing, which has high throughput and low error rates. Sequencing and library preparation costs are dropping rapidly, whether for reduced-representation methods like restriction-site-associated-DNA sequencing strategy (RAD-seq), or whole-genome sequencing^5,6^. On the other hand, user-friendly tools for the analysis of sequencing data are only starting to become available^3^. Here we present SNPmatch, a simple tool for efficiently identifying the most likely strain from a database of *A. thaliana* strain based on a likelihood model for the given markers (SNPs) in each sample. We validated SNPmatch using the sequenced 1135 genomes of *A. thaliana*.

SNPmatch readily identified correct genotypes with only a few thousand random SNP markers — many fewer than expected from any sequencing effort. We then used SNPmatch to perform a quality check of a lab seed stock collection comprising most of the “1001 Genomes”^3^ and RegMap^7^ panels — a total of ~2000 strains. We performed inexpensive (12$/sample) low-coverage sequencing of the seed stock collection by multi-plexing 192 libraries into a single Illumina sequencing lane, resulting in a median sequencing coverage of 1.8X per sample. Using SNPmatch, a staggering 10% of the strains were found to be incorrect, either due to mix-ups within the collection, or with unknown strains. These mistakes are currently being investigated and remedied. This clearly demonstrates the need for and utility of quality control by sequencing.

SNPmatch is a Python library which can be run on the command line, and we have also developed AraGeno (<u>https://arageno.gmi.oeaw.ac.at</u>), a simple web interface that allows user to query their own SNP data against the public polymorphism databases.

## Methods

### Sequencing preparation

3-5 gas-sterilised seeds of each *Arabidopsis* strain were sown directly on soil and stratified for 6 days at 4ºC in the dark. Plants were grown at 10ºC under long days (16 h light, 8 h dark) with 60 % humidity. DNA from 2-3 leaves was extracted using the NucleoMag Plant Kit (Macherey Nagel, cat. # 744400). During the procedure of verifying our lab seed stock collection we modified the library preparation protocol as described below (Fig. 1a).

**Figure 1.**
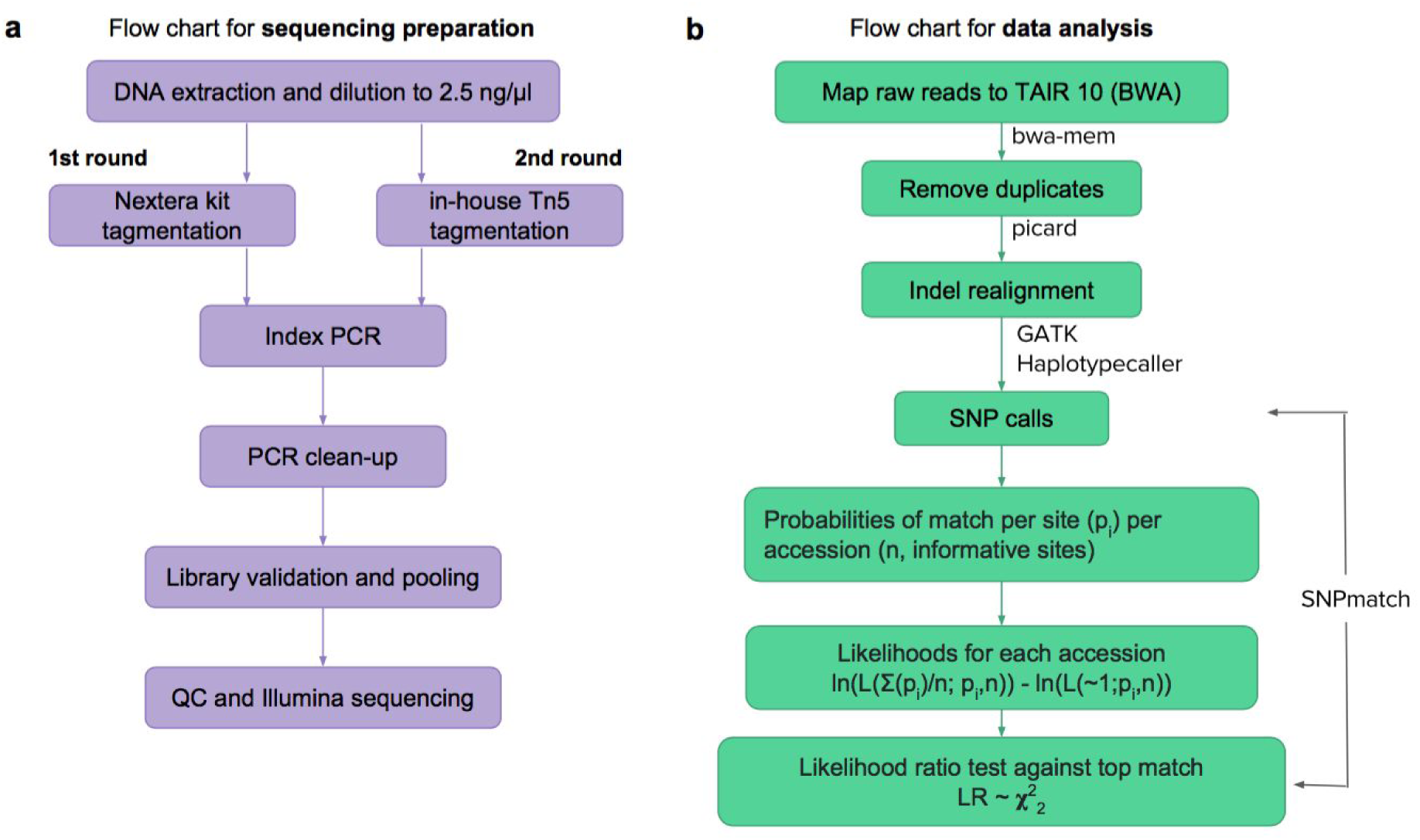
Schematic of library preparation (a) and data analysis (b) workflow.

For the first round of sequencing (1^st^ replicate of the 1001 Genomes collection + RegMap panel) libraries were prepared as described by Baym^8^. Briefly, 2.5 ng of DNA was tagmented and adapter ligated in a 2.5 ul reaction volume using Illumina Nextera™ Kit. Libraries were amplified by Illumina TrueSeq Primers and VeraSeq High Fidelity DNA Polymerase (Biozym, cat #280390). Size selection and PCR clean-up was performed with Agencourt AMPure Beads (Beckman Coulter, cat. #A63882). Libraries were validated with Fragment Analyzer™ Automated CE System (Advanced Analytical) pooled in equimolar concentration for 96-multiplex. Libraries were sequenced on Illumina HiSeq™ 2500 Analyzers using manufacturer’s standard cluster generation and sequencing protocols in 125 bp PE mode.

For the second round of sequencing (2^nd^ replicate of the 1001 Genomes collection) we used a Tn5 transposase, which was generated in-house according to the method described by Picelli^9^. For transposome assembly 0.143 vol of a 100 uM equimolar mixture of preannealed Tn5MEDS-A and Tn5MEDS-B adapters was added to a 1:20 in dilution buffer (50 mM Tris-HCl pH 7.5, 100 mM NaCl, 0.1 mM EDTA, 50 % glycerol, 0.1 % Triton X-100, 1 mM DTT) diluted Tn5 stock and incubated for 30 min at 37ºC. Assembled Tn5 was stored at -20 degrees and used for up to two months. Newly assembled Tn5 was tested on different DNA concentrations each time before usage. Tn5 reactions were assembled in small volumes (2.5 ul total) by mixing 0.5 ul TAPS buffer (50 mM TAPS-NaOH pH 8.5, 25 mM MgCl2, 50 % DMF), 0.1 ul of preassembled Tn5, 1.0 ul DNA (2.5 ng/ul) and 0.9 ul H2O. Reactions were incubated for 7 minutes at 55ºC. Tagmented DNA was used directly for PCR amplifications. PCR was performed by mixing 2.5 ul of tagmented DNA with 11.2 ul VeraSeq High Fidelity DNA Polymerase (Biozym, cat #280390) and 8.8 ul of 0.5 uM index oligos. The PCR program was as follow: 3 min 72ºC, 5 min 98ºC, 8-14 cycles of 10 sec 98ºC, 30 sec 63ºC, 30 sec 72ºC, and 5 min 72ºC. PCR reactions were purified by adding 23 ul AMPure XP beads (cat. # A63882, Beckman Coulter) to 22.5 ul PCR volume. DNA was eluted in 25 ul H_2_O. Libraries were validated with Fragment Analyzer™ Automated CE System (Advanced Analytical) pooled in equimolar concentration for 192-multiplex. Libraries were sequenced on Illumina HiSeq™ 2500 Analyzers using manufacturer’s standard cluster generation and sequencing protocols in 50 bp PE mode.

The modified sequencing protocol of second round were validated by comparing SNPmatch results with that of first round sequencing (Fig. 2). Even with a lower coverage for the samples in second round due to the higher multi-plexing, they gave an unambiguous match to strains as good as for the samples in first round sequencing, indicating a sufficient coverage and library preparation to genotype the samples using SNPmatch.

**Figure 2.**
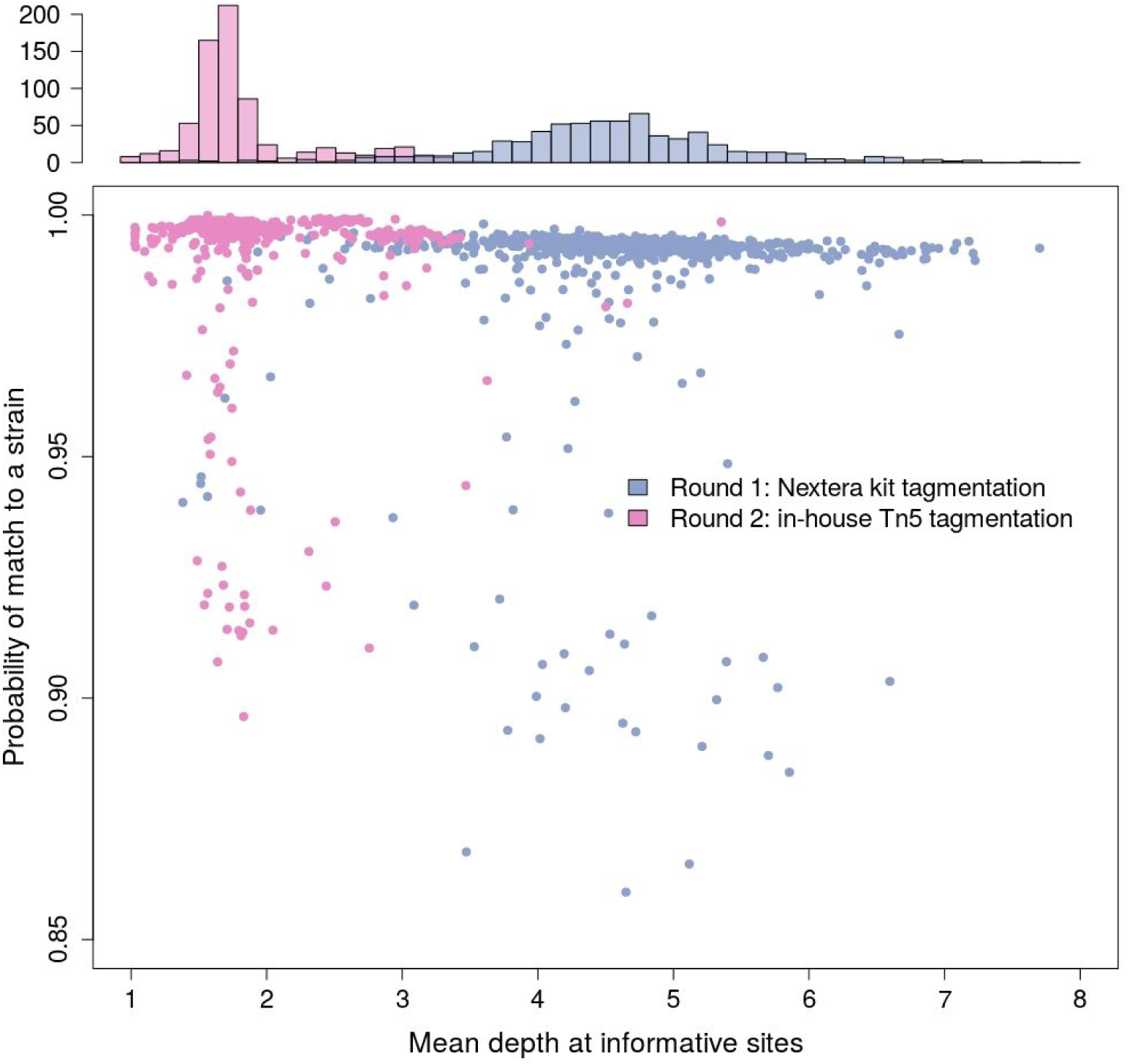
Protocol validation. 1^st^ round of sequencing by Illumina Nextera/96-plex 125 bp paired-end (purple) and 2^nd^ round of sequencing by in-house Tn5/192-plex 50 bp paired-end (pink).

**Figure 3.**
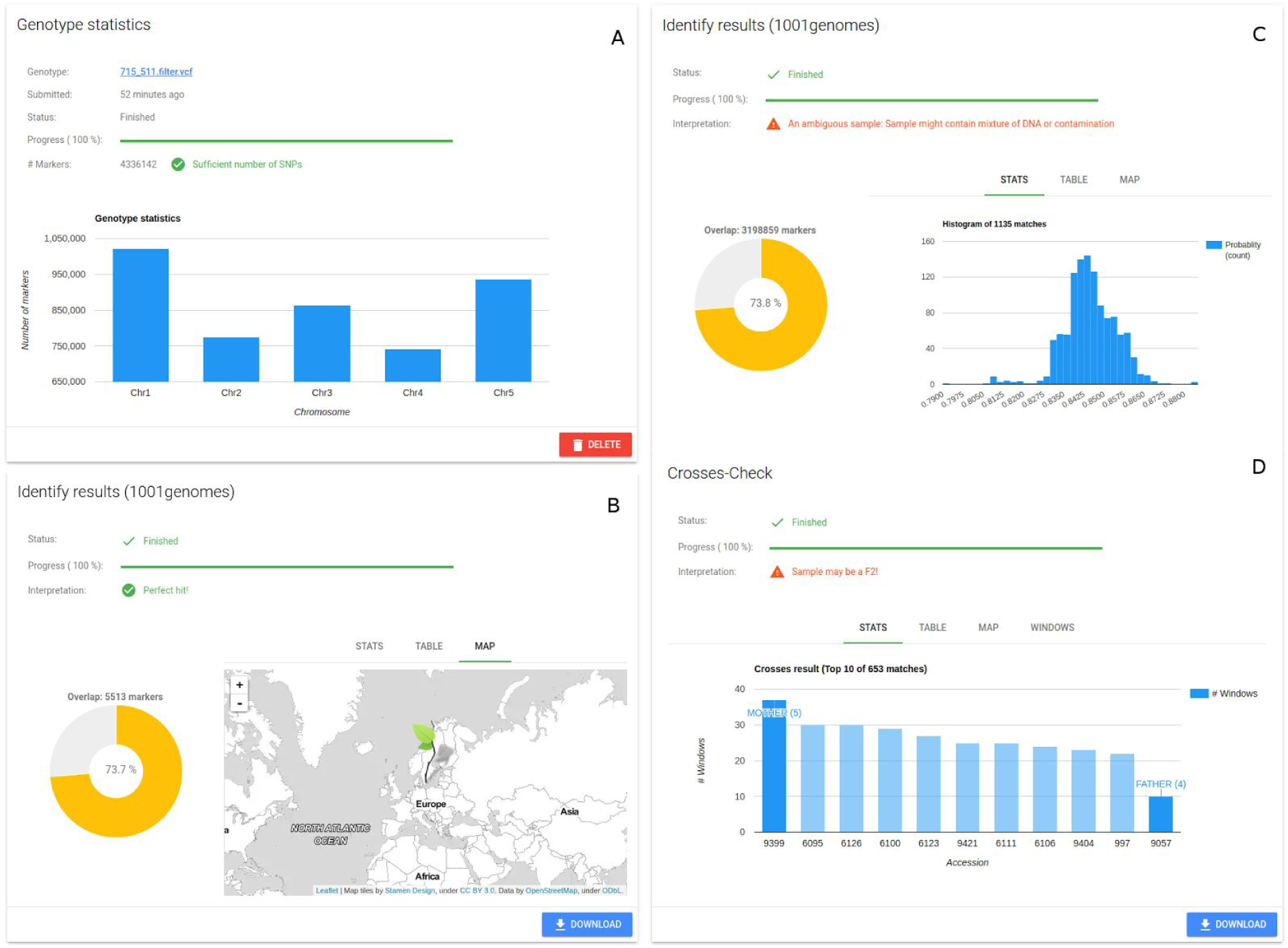
AraGeno web-application. (**A**) General information about the uploaded genotype (VCF or BED) as well as the status of the parsing is displayed. For each supported SNP matrix (Regmap and 1001) the results of the SNPmatch analysis is displayed in separate cards (**B**, **C**). The overlap between the user-provided genotype and the SNP matrix is displayed using a pie-chart. Depending on the analysis result either a histogram or a column chart is displayed of the top matches. Furthermore a table and map provide additional information about the highest matched accessions. If SNPmatch detects that the sample might be a potential cross, an additional analysis will be automatically started and in case of a cross information about the parents are displayed in a separate panel (**D**)

### Sequencing data analysis

All sequencing reads for all the samples were processed accordingly to a standard pipeline, outlined in Fig. 1. In brief, reads were aligned to the *A. thaliana* reference genome (TAIR10, <u>https://www.arabidopsis.org/index.jsp</u>) using bwa mem^10^ with default parameters. Next, duplicates were marked and removed using picard tools software suite (<u>https://broadinstitute.github.io/picard/</u>) using default parameters. SNP and indel calls were obtained from alignment files using GATK HaplotypeCaller^11^ following precisely ‘Best Practice’ instructions on joint genotyping for all the samples per plate. Finally, we filtered for biallelic SNPs using “Select Variants” of GATK.

### SNPmatch algorithm

The input to SNPmatch are SNPs in variant calling format file (VCF file). SNPmatch then performs following calculations to identify the most likely accession from a database.

i. Calculate probability for a match to each accession (a) in the database. This is calculated using the genotype probability scores (PL) from GATK. 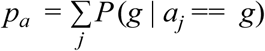

where, j is the SNP position and a_j_ is the genotype of accession A at position j
ii. Likelihoods are calculated for each accession based on the binomial distribution.

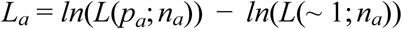

where,
n_a_ is the number of informative sites between a and sample.
iii. Likelihood ratios (LR) are calculated for each accession with the top likely accession.

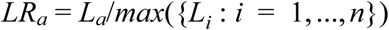

Under the assumption that the LR is chi-squared distributed, a threshold of 3.841 gives a list of accessions that are not significantly different from the top hit (at 95% confidence level).

### AraGeno

SNPmatch is a Python library and primarily used as a command line tool. We made sure that the installation of SNPmatch is as simple and straightforward as possible by hosting SNPmatch on the official Python software repository PyPi (<u>https://pypi.python.org/pypi</u>). Nevertheless we wanted to go one step further and provide SNPmatch as a service to the Arabidopsis community and thus taking care of the installation process and data handling. To this end we developed AraGeno (<u>https://arageno.gmi.oeaw.ac.at</u>), a web-application that exposes a web based frontend to SNPmatch. Users only have to provide a VCF or BED file and AraGeno will take care of validating the data, running SNPmatch on our High Performance Cluster (HPC) and displaying the results of the analysis in an interactive web-interface.

Users can download the results and run as many analysis as they like. By using our HPC system we can easily process many analyzes concurrently. Furthermore AraGeno also provides a RESTful API for programmatic access to the SNPmatch pipeline. AraGeno is hosted on our on-premises cloud infrastructure.

It is a Python based web-application using the Django framework (https://www.djangoproject.com/) and Django REST framework (http://www.django-rest-framework.org/) for the backend and Vue.js (https://vuejs.org/) for the interactive front end side.

In the future we plan to extend the service to additionally allow researchers to mail us samples for genotyping and identifying.

## Code availability

Both the source code for SNPmatch and AraGeno are hosted in GitHub (<u>https://github.com/Gregor-Mendel-Institute/SNPmatch</u>, <u>https://github.com/Gregor-Mendel-Institute/AraGeno</u>).

## Data Records

Resequencing data of all the lines are available on NCBI SRA database with Bioproject accession number PRJNA374784.

## Technical Validation

We validated SNPmatch using the raw data from published 1001 genomes of *A. thaliana*^*3*^, thinning the data to one, three and six million reads for each sample to test the effect of coverage (one million reads roughly correspond to multiplexing 192 *A. thaliana* samples in a single Illumina Hi-Seq 2500 lane, or roughly 1 x coverage). The results were essentially unaffected by this level of thinning, indicating that even 192-fold multiplexing yielded more than enough SNPs to distinguish the strains.

To investigate the required number of SNPs further, we randomly selected subset of SNPs, and ran SNPmatch using those subset only. As illustrated in Fig. 4, the number of strains identified uniquely is largely independent of the number of SNPs, provided that number exceeds a few thousand — as long as we do not try to distinguish very closely related strains. In the 1001 genomes panel, there are 78 North American strains and 40 other pairs of strains that differ by fewer than 1k SNPs, and there are 60 pairs of strains that differ by less than 50k SNPs^12^. These strains are difficult to distinguish from each other even with millions of SNPs.

**Figure 4.**
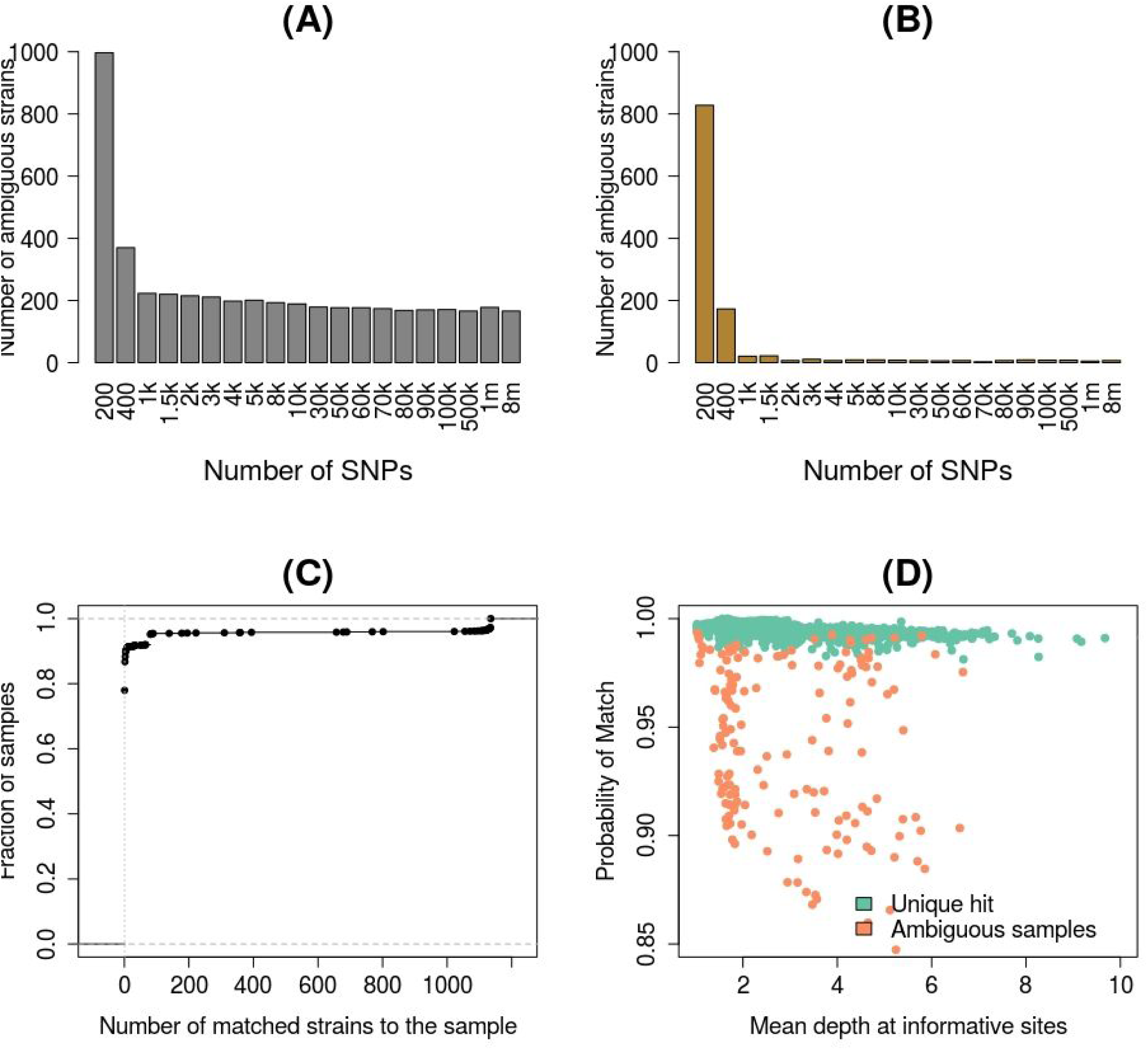
Validation of SNPmatch and 1001 seed stock collection. **(A, B)** Number of ambiguous strains resulting with markers (SNPs) randomly sampled from entire (A) and collapsed (B) 1001 SNP matrix respectively **(C, D)** Genotyping results for the samples resequenced from 1001 seed stock collection. Samples are identified unambiguously to a strain even with low-sequencing coverage.

Next, we used SNPmatch to validate our own seed stocks. We resequenced almost complete sets of the “1001 Genomes”^3^ and “RegMap”^7^ collections using the protocols described in Methods. Of a total of 1998 sequenced strains, 1991 yielded sufficiently good SNP data for analysis. Of these, 1797 were assigned to the correct strain (or set of strains differing by less than 5000 SNPs in the databases). Of the remaining 194 strains, 82 (30 of the “1001 Genomes” and 52 of the “RegMap” collections) were unambiguously assigned to the wrong strain (or set of closely related strains), indicating sample or strain mix-up (suppl. Table 1). Finally, the remaining 112 (44 of the “1001 Genomes” and 68 of the “RegMap” collections) did not match any strain in the databases (suppl. Table 1). These samples could represent unknown strains, DNA contamination, or outcrossed individuals, however DNA contamination and outcrossing should both result in an unusually large number of heterozygous calls, which we do not observe (Fig. 5). Therefore, we conclude that the ambiguous calls are most likely due to sample mix-up with unknown strains. We are currently collecting and verifying the 194 incorrect strains from different sources in order to make sure that the stock center has the right germplasm.

**Figure 5.**
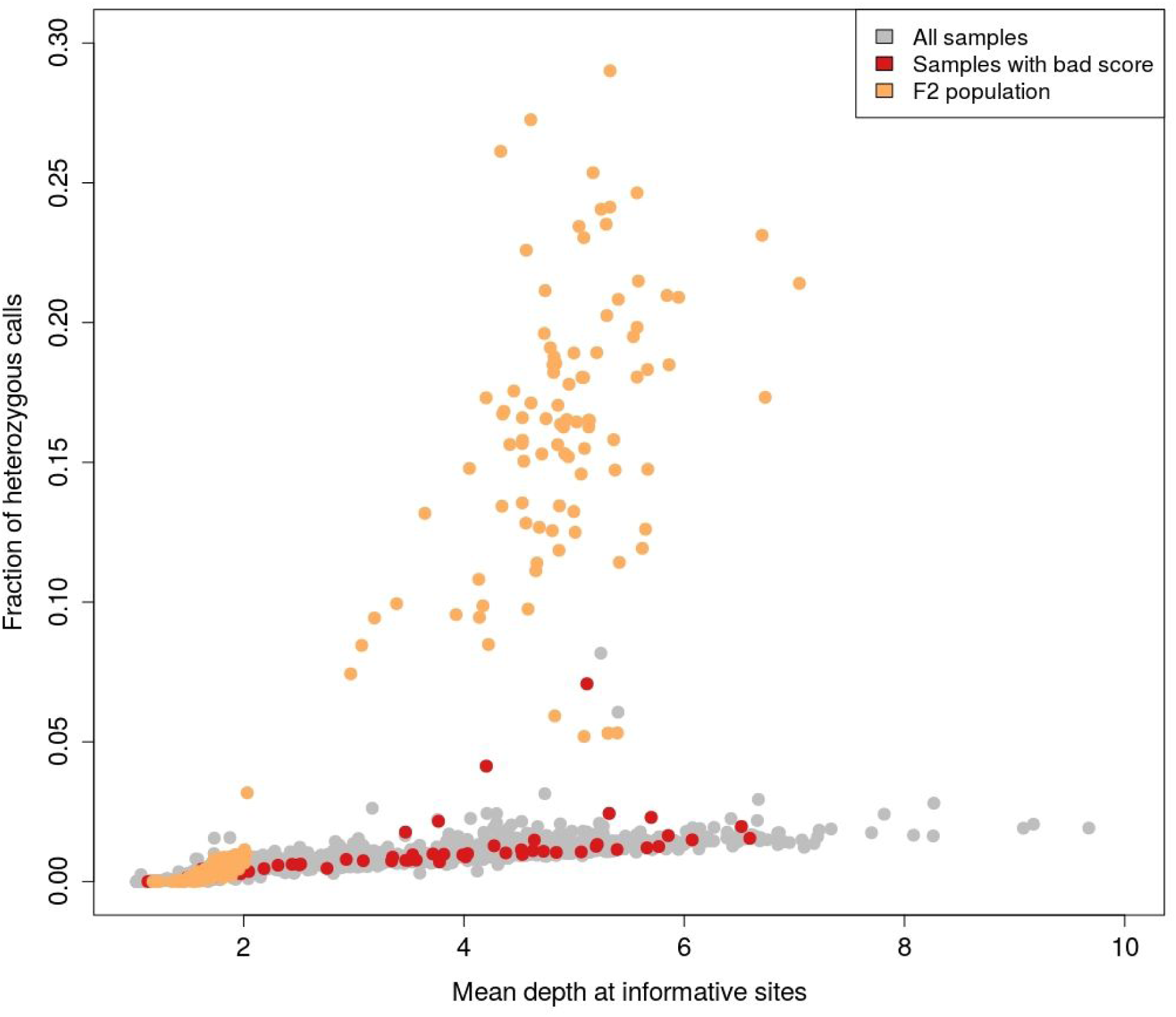
Fraction of heterozygous calls in the samples which do not unambiguously match to any strain (red) are comparable to all the other samples. Whereas F2 population (orange) have much higher fraction of heterozygous calls when their coverage is good.

At least when the parent strains are in the database, SNPmatch can be used to identify hybrid individuals. Such individuals will not find an unambiguous match in the database, but will be equally closely related to both parents (Fig. 6A). For such individuals running SNPmatch in windows across the genome will quickly reveal its hybrid nature (Fig. 6B).

**Figure 6.**
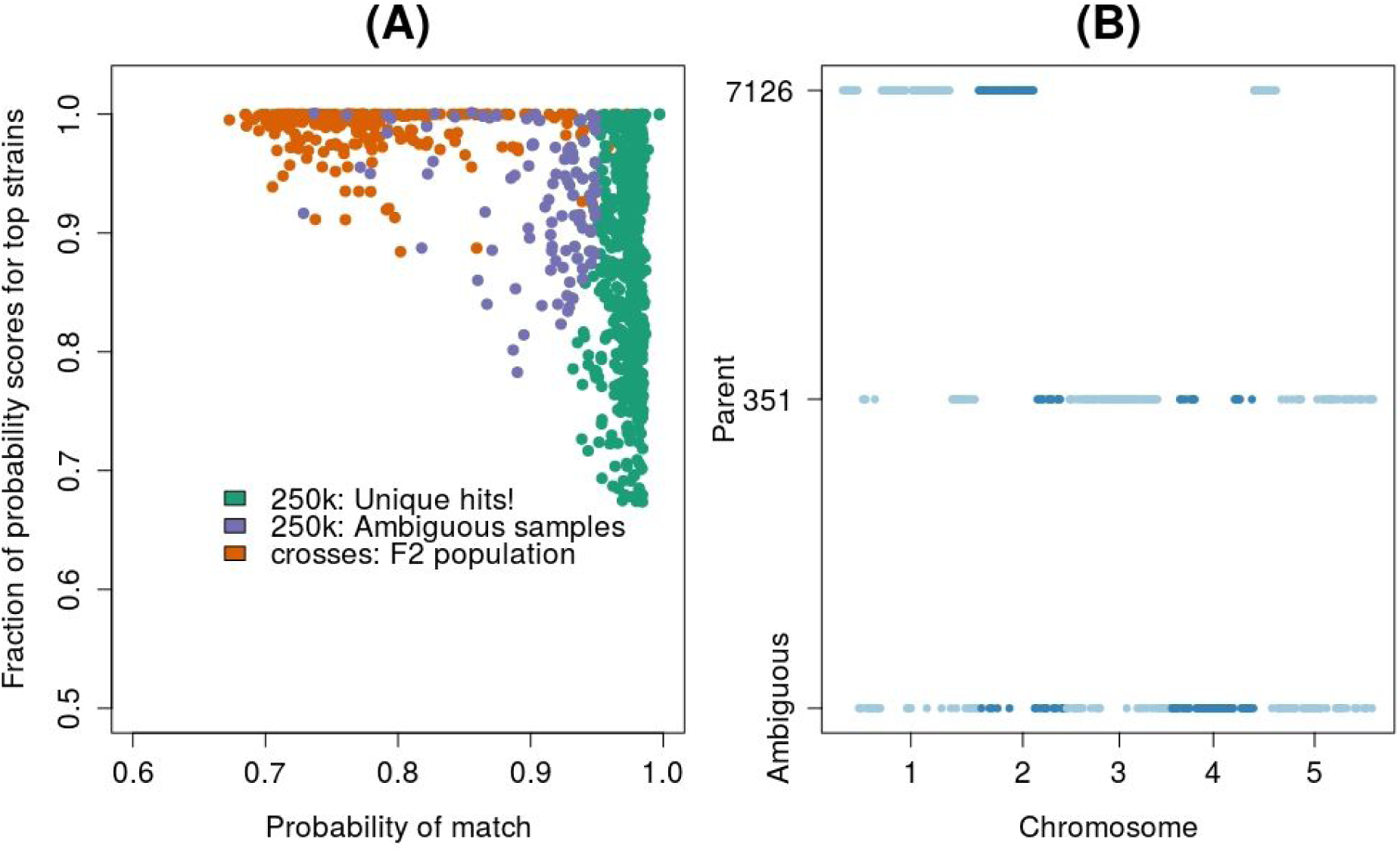
Identification of the parents in a F2 population using SNPmatch and validation of 250k seed stock.

## Acknowledgements

We thank members of Nordborg lab for the discussions and VBCF NGS unit for performing the Illumina sequencing.

